# A versatile human intestinal organoid-derived epithelial monolayer model for the study of enteric pathogens

**DOI:** 10.1101/2020.11.24.397141

**Authors:** Kourtney P. Nickerson, Alejandro Llanos-Chea, Laura Ingano, Gloria Serena, Alba Miranda-Ribera, Meryl Perlman, Rosiane Lima, Marcelo B. Sztein, Alessio Fasano, Stefania Senger, Christina S. Faherty

**Affiliations:** Mucosal Immunology and Biology Research Center, Division of Pediatric Gastroenterology and Nutrition, Massachusetts General Hospital, University of Maryland School of Medicine, Baltimore, Maryland; Department of Pediatrics, Harvard Medical School, Boston, Massachusetts, University of Maryland School of Medicine, Baltimore, Maryland; Center for Vaccine Development and Global Health, Department of Pediatrics, University of Maryland School of Medicine, Baltimore, Maryland

**Keywords:** *Shigella*, *Salmonella*, *Escherichia coli*, infection models, human, intestinal, organoid, enteroid, HIODEM, DAPT, RANKL, epithelial monolayer, enterocytes, goblet cells, mucus, M cells

## Abstract

Gastrointestinal infections cause significant morbidity and mortality worldwide. The complexity of human biology and limited insights into host-specific infection mechanisms are key barriers to current therapeutic development. Here, we demonstrate that two-dimensional epithelial monolayers derived from human intestinal organoids, combined with *in vivo*-like bacterial culturing conditions, provide significant advancements for the study of enteropathogens. Monolayers from the terminal ileum, cecum, and ascending colon recapitulated the composition of the gastrointestinal epithelium, in which several techniques were used to detect the presence of enterocytes, mucus-producing goblet cells, and other cell types following differentiation. Importantly, the addition of receptor activator of nuclear factor kappa-B ligand (RANKL) increased the presence of M cells, critical antigen-sampling cells often exploited by enteric pathogens. For infections, bacteria were grown under *in vivo*-like conditions known to induce virulence. Overall, interesting patterns of tissue tropism and clinical manifestations were observed. *Shigella flexneri* adhered efficiently to the cecum and colon; however, invasion in the colon was best following RANKL treatment. Both *Salmonella* Typhi and Typhimurium serovars displayed different infection patterns, with *S*. Typhimurium causing more destruction of the terminal ileum and *S*. Typhi infecting the cecum more efficiently than the ileum, particularly with regards to adherence. Finally, various pathovars of *Escherichia coli* validated the model by confirming only adherence was observed with these strains. This work demonstrates that the combination of human-derived tissue with targeted bacterial growth conditions enables powerful analyses of human-specific infections that could lead to important insights into pathogenesis and accelerate future vaccine development.

**Importance:** While traditional laboratory techniques and animal models have provided valuable knowledge in discerning virulence mechanisms of enteric pathogens, the complexity of the human gastrointestinal tract has hindered our understanding of physiologically relevant, human-specific interactions; and thus, has significantly delayed successful vaccine development. The human intestinal organoid-derived epithelial monolayer (HIODEM) model closely recapitulates the diverse cell populations of the intestine, allowing for the study of human-specific infections. Differentiation conditions permit the expansion of various cell populations, including M cells that are vital to immune recognition and the establishment of infection by some bacteria. We provide details of reproducible culture methods and infection conditions for the analyses of *Shigella, Salmonella*, and pathogenic *Escherichia coli* in which tissue tropism and pathogen-specific infection patterns were detected. This system will be vital for future studies that explore infection conditions, health status, or epigenetic differences; and will serve as a novel screening platform for therapeutic development.

## Introduction

Over 10 million pediatric deaths occur annually, with over half of these deaths resulting from microbial infection (1). Despite improvements in hygiene, availability of clean water, and access to treatment, few preventive therapies exist and pediatric mortality rates remain high (1-5). Gastrointestinal (GI) pathogens such as *Shigella, Salmonella*, and *Escherichia coli* cause diarrhea, dysentery, and even sepsis in some instances (6-10). Many of these pathogens are human-restricted, rendering traditional laboratory models insufficient for understanding disease pathologies in humans. In fact, the complexities of human biology coupled with human-specific infection patterns are key barriers to understanding pathogenic mechanisms for successful vaccine development.

To overcome these barriers, we further developed methodology to transition human intestinal stem cells from spheroid cultures into single two-dimensional (2-D) cell monolayers that recapitulate the various cell types of the GI tract (11) and answer critical questions regarding how human-specific pathogens interact with host cells. Herein, we present detailed protocols to prepare human intestinal organoid-derived epithelial monolayers (HIODEM) isolated from the ileum, cecum, and ascending colon to test infection with *Shigella, Salmonella*, and pathogenic *E. coli*. Monolayers were composed of enterocytes, goblet cells, and other cell types of the various intestinal segments. Through ligand stimulation, the monolayer system was modified to promote expansion of the microfold (M) cell population of the follicle-associated epithelium often exploited by bacterial pathogens to gain access to the epithelium (12, 13). Both adherence and invasion of the three GI segment-derived models were examined with the various pathogens, and the role of M cells in enteric infections was evaluated for *Shigella* and *Salmonella*. We also utilized bacterial growth conditions that replicate the host environment to enhance the virulence of the pathogens prior to infection. In all, this combined methodology has significant potential to transition current research methods into the most human-specific, *in vivo*-like model to study bacterial pathogenesis and preclinically evaluate therapeutic candidates, potentially leading to paradigm-shifting approaches to vaccine or therapeutic development.

## Results

### The HIODEM model for studying enteric pathogenesis

Organoids were derived from tissue biopsies collected from the terminal ileum, cecum, and ascending colon, and subcultured for monolayer generation. After isolation and propagation, crypt stem cells were dissociated into single cells and seeded onto transwell inserts. Monolayers reached confluency in 7 to 10 days, in which transepithelial electrical resistance (TEER) was monitored to indicate formation of functional barriers (14). TEER readings were the highest in the terminal ileum averaging 1010 +/- 112 Ω*cm^2^ with 0.33 cm^2^ transwells, whereas cecum monolayers averaged 490 +/- 31 Ω*cm^2^ and the colon had much lower TEER values averaging at 209 +/- 2.7 Ω*cm^2^ (**Table 1**). Following the stabilization of TEER readings, monolayers were treated with the γ-secretase inhibitor N-[N-(3,5-Difluorophenacetyl)-L-alanyl]-S-phenylglycine t-butyl ester (DAPT) (11, 15), with or without the receptor activator of nuclear factor kappa-B ligand (RANKL) (16) at physiological (100 ng/ml) or supraphysiological (500 ng/ml) doses for 24 or 48 hours to ensure differentiation and provide the immune stimulation to induce M cell expression (16-18).

**Table 1.**
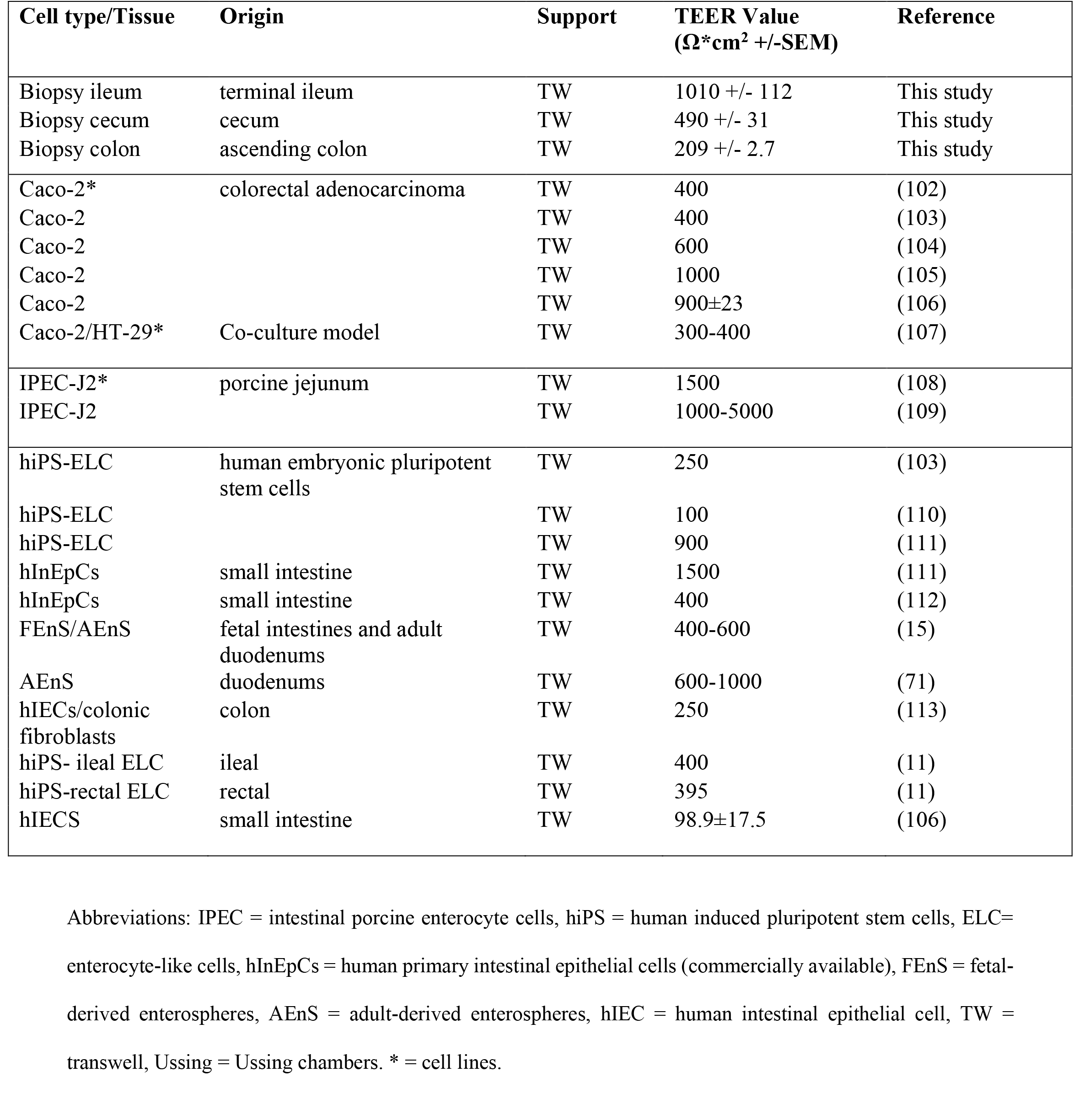
Average TEER values from this study and reported in the literature.

Several methodologies were used to characterize the differentiated monolayers, including flow cytometry, reverse transcription qPCR (RT-qPCR), and transmission electron microscopy (TEM). We focused on ensuring the presence of enterocytes, mucus-producing goblet cells, and M cells, with other cell markers evaluated by RT-qPCR only. First, for flow cytometry, mature monolayers were examined using antibodies against the transcription factors ESE1 for enterocytes (19), KLF4 for goblet cells (20), and SPIB for M cells (17). The analyses confirmed the presence of the respective cellular markers and enabled us to estimate the cellular populations for each cell type (**Supplemental Figure S1**). Specifically, for the M cell phenotype, RANKL treatment increased the percentage of SPIB^+^ cells relative to the DAPT only treatment across all tissue types with both doses of RANKL for 24 and 48 hours (**Figure 1**). Second, to confirm the induction of the *SPIB* gene with RANKL treatment, we analyzed gene expression at 24 hours following 100 ng/ml RANKL treatment (**Figure 2**). The RT-qPCR analyses revealed significant induction of *SPIB* expression with the DAPT + RANKL treatment relative to the DAPT only treatment, with no or minor changes in expression for *ESE1, KLF4*, and other cellular markers (**Figure 2** and **Table 2**). It is important to note that expression of *ESE1, KLF4*, and the other genes was detected in all conditions (data not shown), but the changes in expression were not significantly altered in DAPT + RANKL treatment compared to DAPT only treatment. Finally, mature monolayers were evaluated by TEM to visualize and confirm the various cell types, monolayer quality, and barrier formation (**Figure 3**). Characteristic features of enterocytes, goblet cells, and M cells were visualized, which included the presence of microvilli for enterocytes, secretory granules for goblet cells, and the absence of microvilli for M cells (21-23). In all, the data demonstrated that multiple cellular populations were present in the HIODEM models evaluated under the differentiation conditions described above and that RANKL treatment increased the presence of M cells.

**Figure 1.**
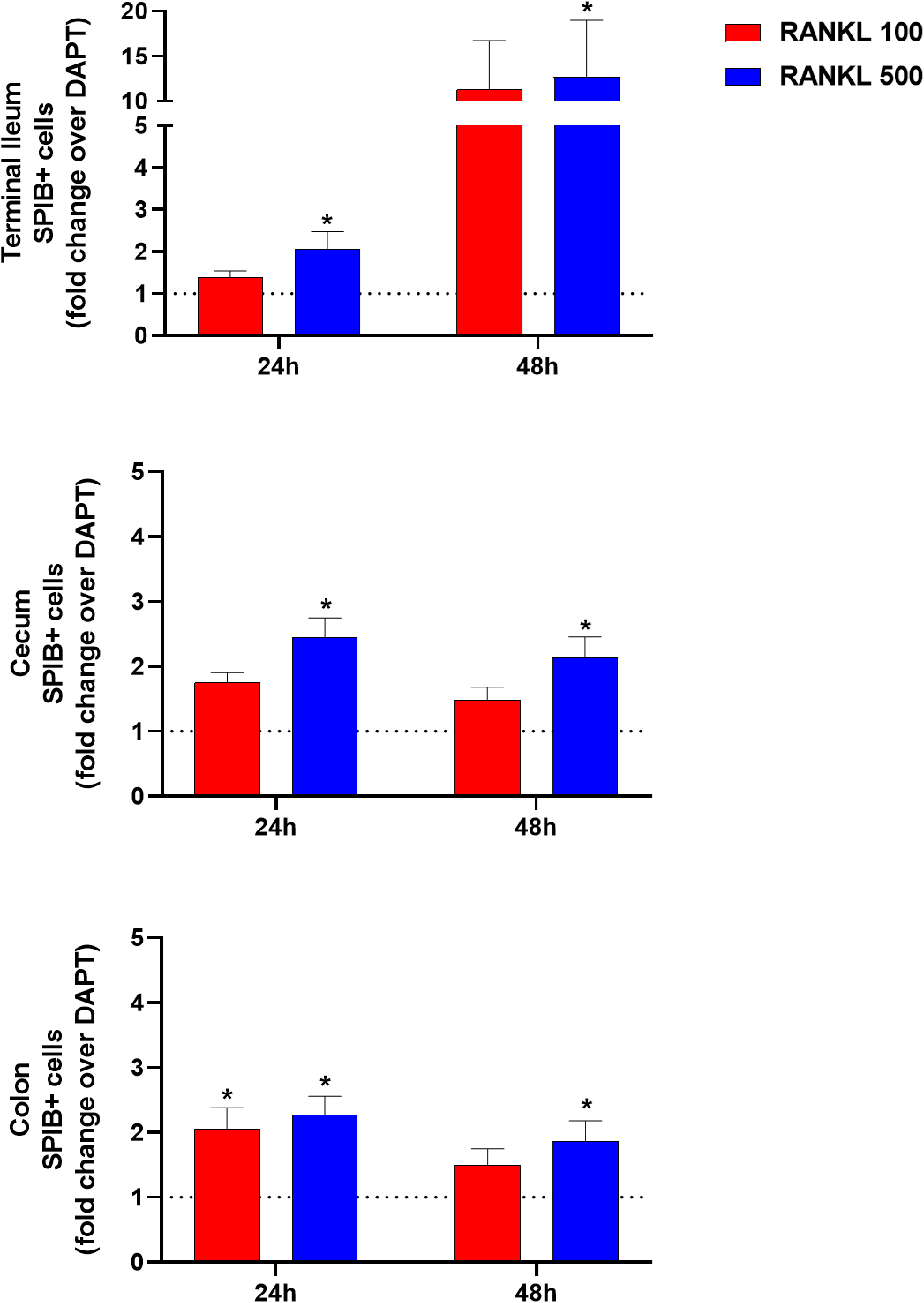
Flow cytometry analyses identify an increase in SPIB^+^ expression following RANKL treatment. The increase in SPIB^+^ expression, which is indicative of M cell expression, after 24 or 48 hours of RANKL treatment is represented as fold change of SPIB^+^ cells in tissues treated with RANKL (either DAPT + RANKL 100ng/ml, red bars or DAPT + RANKL 500 ng/ml, blue bars) over the matching tissues treated only with DAPT (represented by the dotted line) for each tissue. Statistical significance was determined by paired Friedman test for the DAPT + RANKL (100 or 500 ng/ml) differentiation compared to DAPT only-differentiation of the matched originating tissue (*, p <0.05). A total of 4 biological replicate experiments were analyzed for the terminal ileum and cecum, and 6 biological replicates were analyzed for the ascending colon. For each biological sample, an entire 12- well plate was trypsinized, and the cells were pooled into one sample tube for staining and analysis.

**Figure 2.**
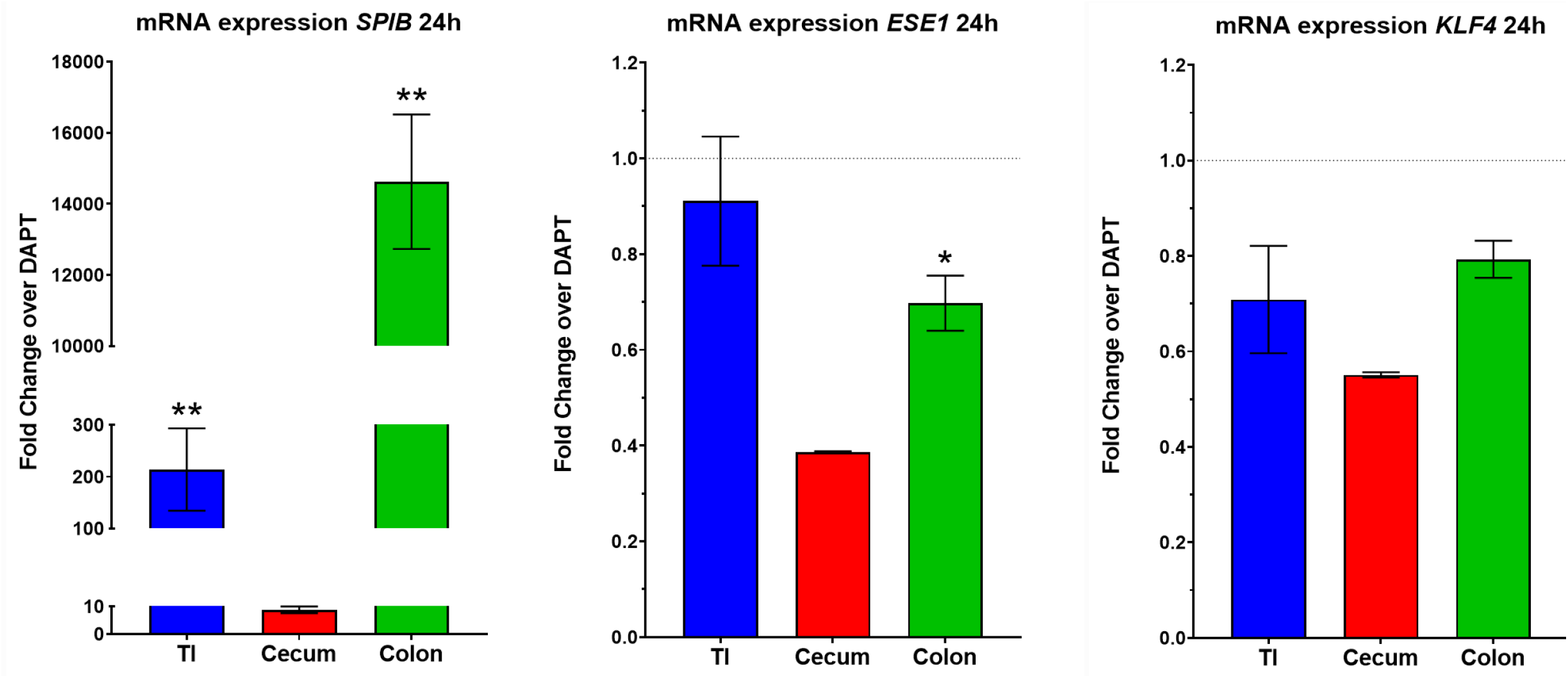
RT-qPCR analyses demonstrate significantly induced *SPIB* gene expression following RANKL treatment. Gene expression for *SPIB, ESE1*, and *KLF4* was measured following 24-hour DAPT + 100 ng/ml RANKL or DAPT only treatment for the terminal ileum (TI, blue, n = 6), cecum (red, n = 3), and colon (green, n= 6). Data are expressed as DAPT + RANKL fold change over the DAPT only treatment, shown as the average fold change +/- the SEM. All data were normalized to expression from the *18S* housekeeping gene. Statistical significance was determined with the Mann Whitney unpaired t-test of the dCt. Data were considered significant at a p-value <0.05 (* < 0.05, ** < 0.01). Please note the different y-axes for each graph.

**Figure 3.**
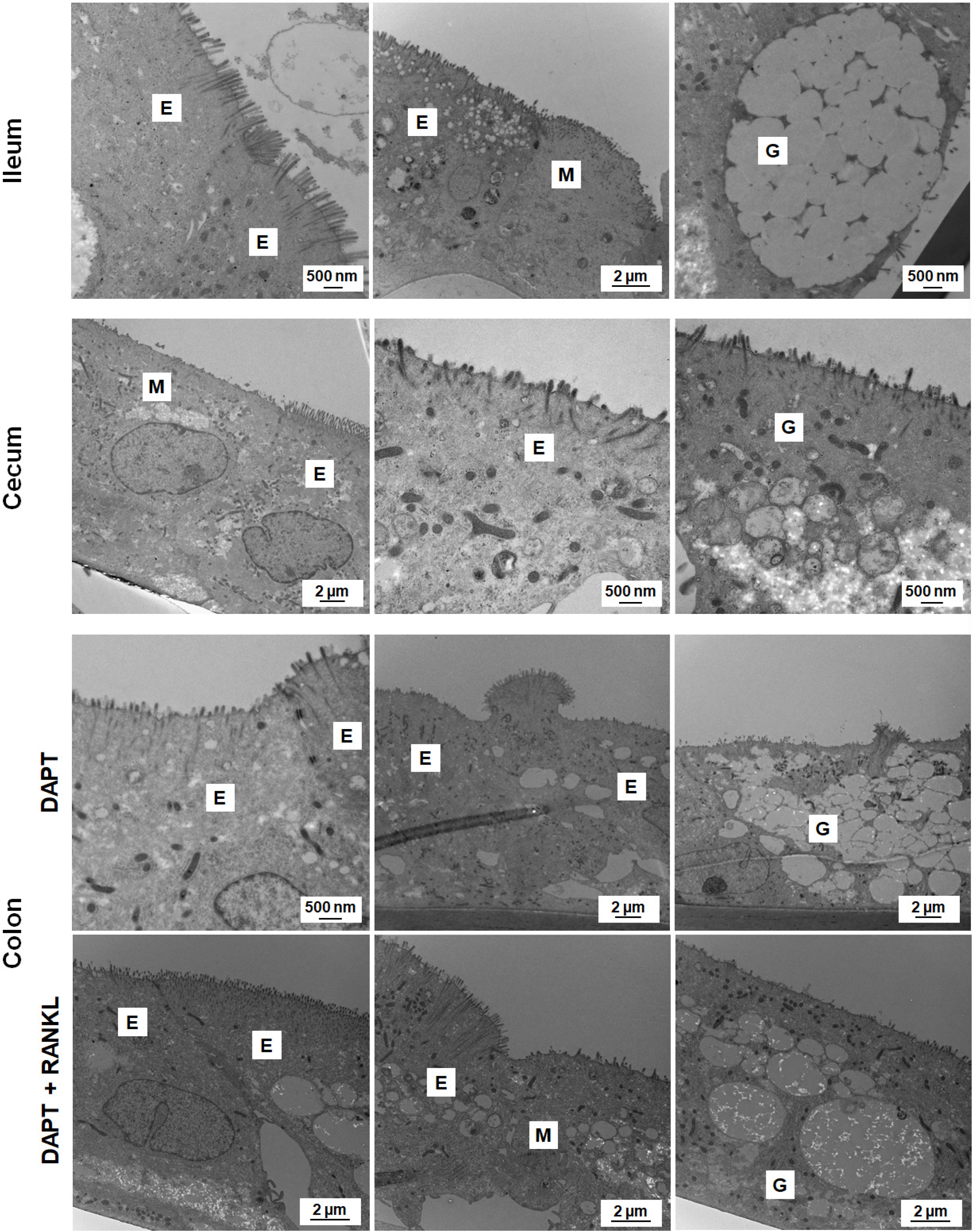
Transmission electron microscopy analysis of the HIODEM models. Mature monolayers derived from the terminal ileum (top), cecum (middle), or colon (bottom) were evaluated by TEM to visualize the cell types present, monolayer quality, and barrier formation. All models displayed characteristic features of enterocytes (E), goblet cells (G), and M cells (M). For the colon, differentiation treatments with DAPT and DAPT + RANKL were compared to visualize the presence of M cells. Magnifications range from 5,000X to 25,000X and the corresponding scale bars are provided.

**Table 2.**
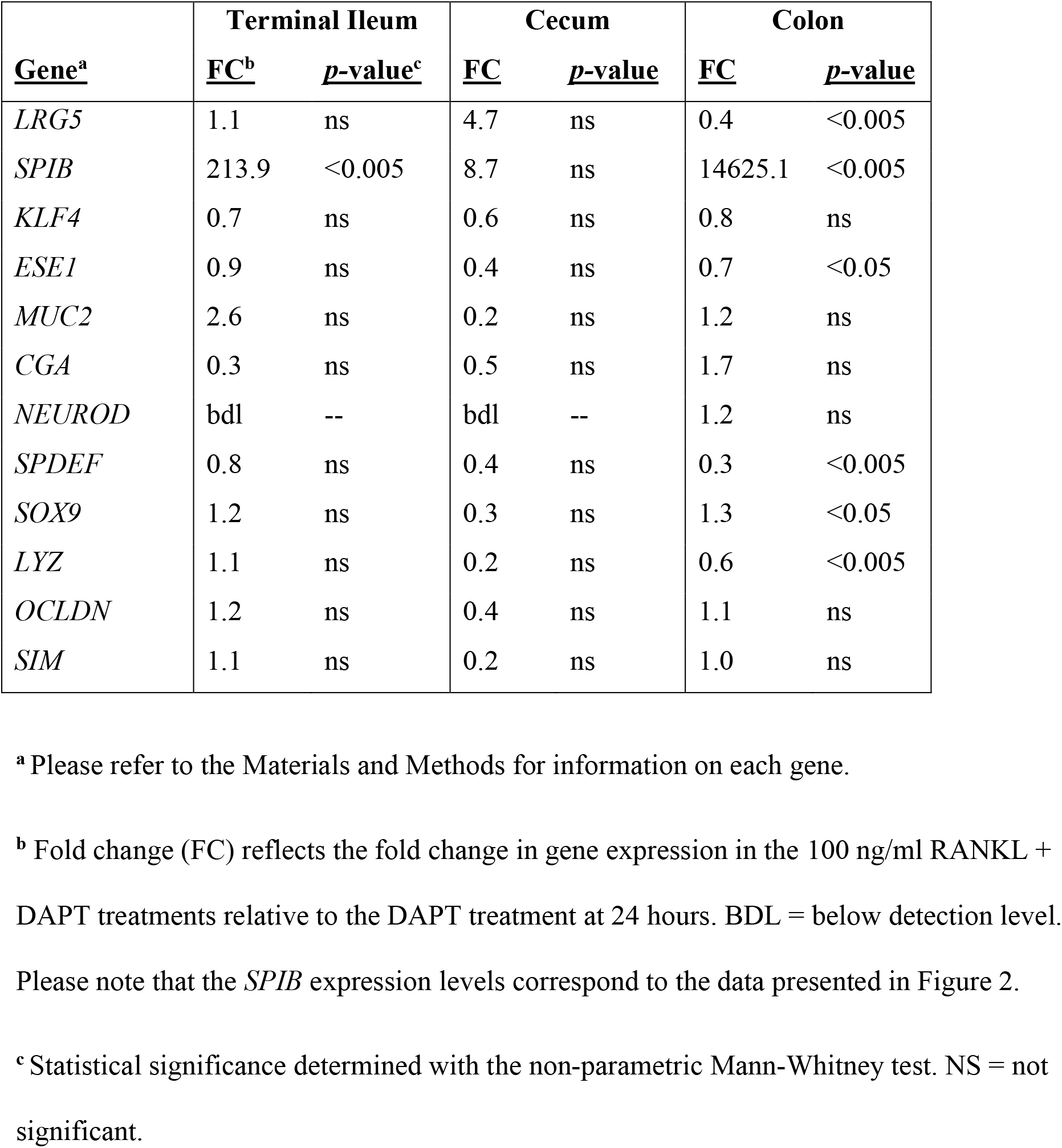
Gene expression analyzed by RT-qPCR in models treated with 100 ng/ml RANK + DAPT for 24 hours.

### Robust *Shigella* infection requires colonic M cells

To validate the site of *Shigella* infection, we analyzed wild type *S. flexneri* strain 2457T infection in the terminal ileum, cecum, and colon-derived HIODEM by assaying for adherence and invasion. Prior to infection, *S. flexneri* was cultured in a combination of bile salts and glucose to replicate small intestinal transit (24-26). For adherence, approximately 5% of the bacterial inoculum adhered to ileum-derived HIODEM, while the adherence rates for cecum and colon-derived HIODEM models were nearly triple (~12-15%; **Figure 4A**). For invasion, rates were consistent across the models under most treatments; however, colon-derived monolayers differentiated with RANKL to promote the maturation of M cells consistently had a three-fold increase in invasion (**Figure 4B**). A virulence plasmid-cured, non-invasive strain of *S. flexneri* (strain BS103 (27)) was used as a negative control for invasion and was unable to invade the models under any conditions. Scanning electron microscopy (SEM) of *S. flexneri* infected colonic models demonstrated attachment to the apical surface of the cells (**Figure 4C**). The *S. flexneri* adherence pattern occurred throughout the surface of the monolayer. Additionally, shadowed areas beneath adherent bacteria appeared on cells lacking microvilli in the RANKL-treated samples, accompanied by perforations in other parts of the cells. These observations appear to be visual representations M cell transit required for *Shigella* invasion (28). Cross-section TEM verified these observations with visualization of invading bacteria localized to M cells (**Figure 4D**). Finally, to further validate the model, *S. flexneri-*infected colon HIODEM monolayers were evaluated for interleukin-8 (IL-8) and lactate dehydrogenase (LDH) release (**Supplemental Figure S2**) since *Shigella* infection is accompanied by IL-8 secretion (29, 30) and inhibition of epithelial cell death (31, 32). Significant IL-8 secretion from *S. flexneri* infection was observed while mock-treated cells had no detectable levels of IL-8. Additionally, very low levels of LDH was detected for *S. flexneri*-infected monolayers with no significant difference to mock infected.

**Figure 4.**
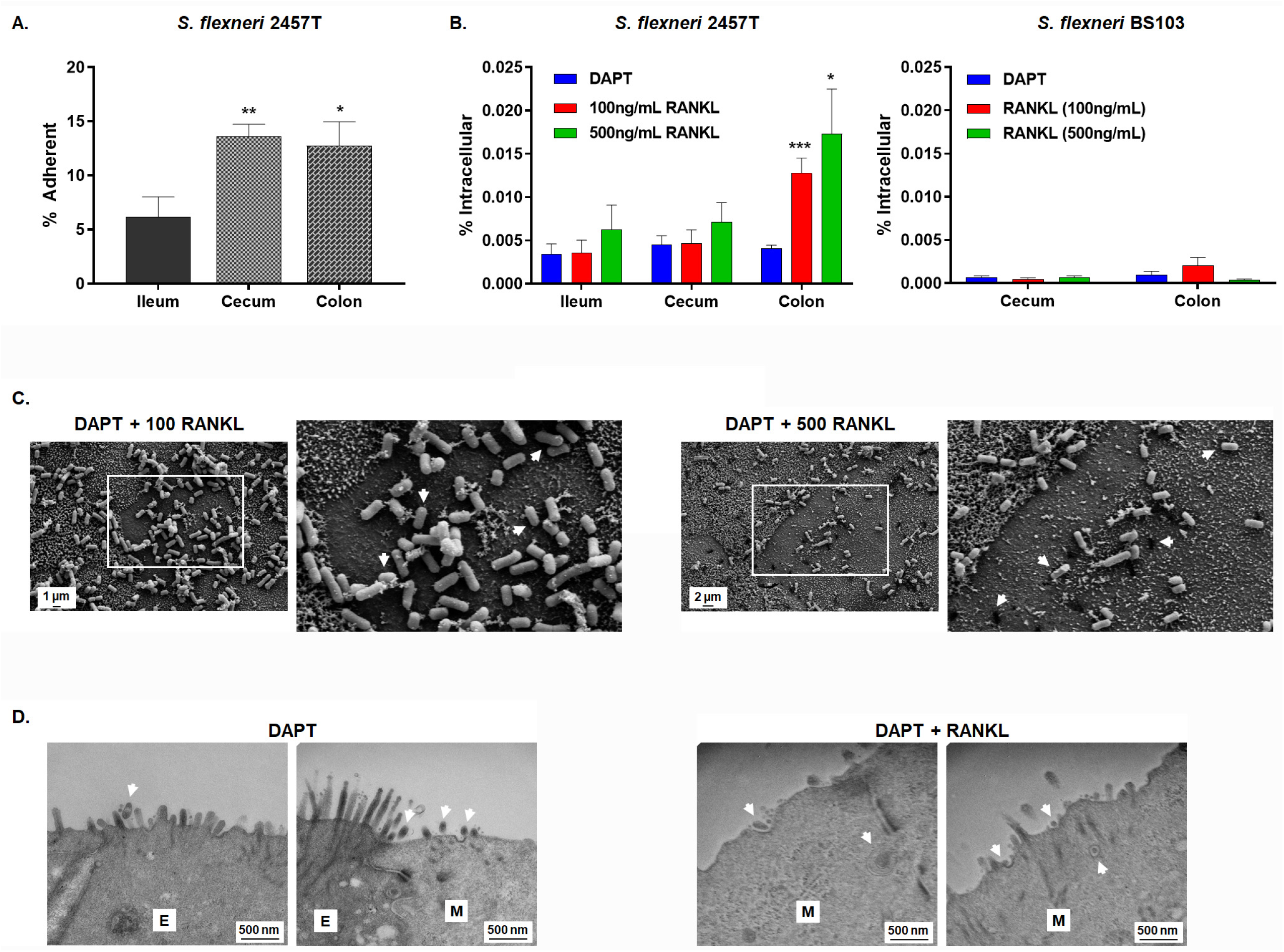
*Shigella* infection analysis reveals a tissue tropism for the colon. Infection analyses were performed with at least three biological replicates, in which three technical replicates were present for each experiment. Imaging analysis was performed on biological or technical independent samples relative to the plotted infection data. For plotted data, please note the different y-axis scales. Statistical significance was determined by the Student’s t-test of the indicated comparisons, in which * = p <0.05, ** = p <0.01, *** = p < 0.001. **A.** Ileum-, cecum-, and colon-derived HIODEM models were infected with *S. flexneri* strain 2457T and the percent of the inoculum adherent to the monolayer surface was determined. *S. flexneri* was adherent to all three tissue locations with a significant increase in adherence to the cecum- and colon-derived HIODEMs relative to ileum-derived HIODEMs. Statistical analyses compared the ileum adherence rates to the cecum or colon. **B.** *S. flexneri* invasion analysis of the HIODEM models. Colon-derived HIODEMs differentiated with DAPT and RANKL had nearly three times as many intracellular *S. flexneri* strain 2457T in colonic RANKL-treated HIODEMs. Statistical analyses compared the DAPT-treated monolayers relative to the two DAPT + RANKL treatments for each set of models (ileum, cecum, and colon). The non-invasive strain BS103 did not have significant recovery titers following gentamicin treatment in the cecum or colon HIODEMs despite M cell differentiation, which validated the invasion data for 2457T. **C.** SEM of colon-derived HIODEMs differentiated with DAPT + RANKL at either 100 ng/ml or 500 ng/ml concentrations. Bacterial adherence to both enterocytes and M cells, with cell surfaces lacking microvilli, were observed. Magnification ranged from 5,000X to 7,000X, with 2 μm and 1 μm scale bars, respectively. Enlarged portion of the images are denoted by the white boxes, arrows point to areas of various stages of bacterial translocation on the surface of the M cells. Please note that not all areas are highlighted. **D.** Transmission electron micrographs of DAPT and DAPT + RANKL-differentiated colonic monolayers infected with *S. flexneri* reveal bacterial association with the apical surface of enterocytes (E) and M cells (M). Areas with bacterial cells at the cellular surface or translocating through M cells are highlighted by the arrows. Magnification for all images is 50,000X with 500 nm scale bars noted.

### Serovar-specific aspects of *Salmonella* infection are revealed using the HIODEM model

To demonstrate versatility of the HIODEM model, monolayers derived from ileum, cecum, and colon tissue were used to examine adherence and invasion of wild type *Salmonella enterica* serovar Typhi strain Ty2 and serovar Typhimurium strain SL1344. *Salmonella* Typhi infection was more prevalent in the cecum, with a higher rate of adherence and a modest increase in invasion relative to the ileum (**Figure 5A**). Furthermore, infection of ileum HIODEMs treated with RANKL to promote M cell maturation did not result in a significant increase in intracellular bacteria (**Figure 5B**). Transmission electron microscopy of infected HIODEM cells revealed *S*. Typhi associated with the enterocyte surface, remodeled host cytoskeleton, or contained within intracellular vesicles (**Figure 5C**); while bacterial association with secreted mucus was also observed following immunostaining analysis (**Supplemental Figure S3**). Interestingly, infection comparison of the *S*. Typhi to *S*. Typhimurium analyses revealed serovar-specific infection patterns (**Figure 5D** and **5E**). Unlike *S*. Typhi, *S*. Typhimurium infected and replicated robustly within ileum monolayers with significant destruction of the monolayers revealed upon SEM analysis. Like the TEM, the SEM analysis of the *S*. Typhi infected monolayers showed bacteria interacting with enterocytes through bacterial surface structures binding to microvilli, with an overall pattern of surface association to intact monolayers. These results both verify our previous findings for *S*. Typhi infection (33) and demonstrate distinctions in serotype-specific infection patterns along different segments of the human GI tract.

**Figure 5.**
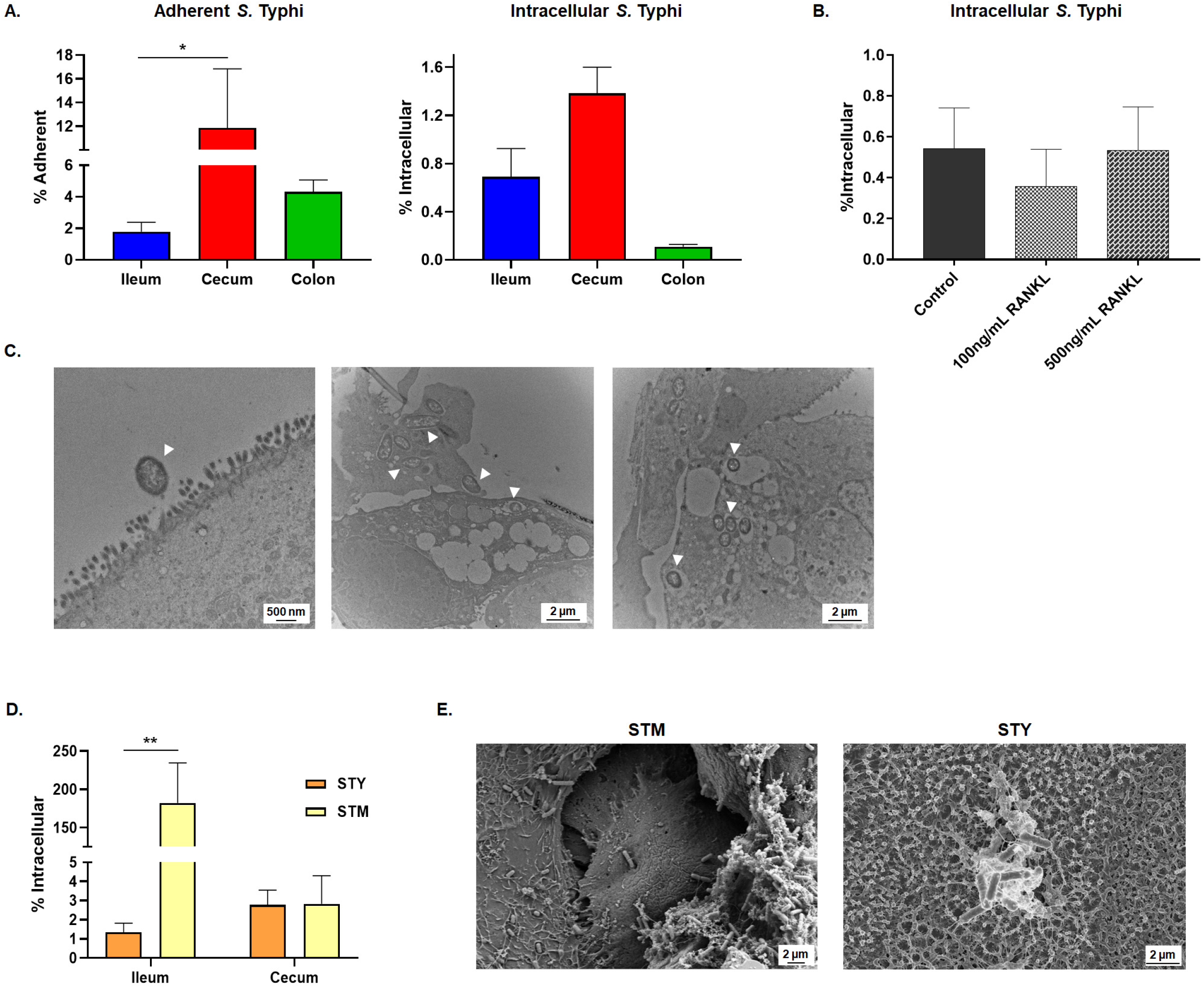
The HIODEM model demonstrates *Salmonella* serovar-specific infection patterns. Infection analyses were performed with at least three biological replicates, in which three technical replicates were present for each experiment. Imaging analysis was performed on biological or technical independent samples relative to the plotted infection data. For plotted data, please note the different y- axis scales. Statistical significance was determined by the Student’s t-test of the indicated comparisons, in which * = p <0.05, ** = p <0.01. **A.** *Salmonella enteric* serovar Typhi strain Ty2 adhered to ileum-, cecum-, and colon-derived HIODEMs, with the greatest rate detected in the cecum monolayers. Invasion was prevalent in the ileum- and cecum-derived HIODEMs, with a modest increase in the cecum monolayers. **B.** RANKL-treatment of the ileum HIODEMs did not result in significant differences in the rate of invasion. **C.** TEM analysis of *S*. Typhi invasion in the ileum HIODEMs. Magnification ranged from 10,000X to 25,000X, with 2 μm and 500 nm scale bars, respectively. Arrows point to bacteria, please note that not all bacterial are highlighted. **D.** *Salmonella enterica* serovar Typhimurium strain SL1344 (STM) preferentially invades ileum HIODEMs at significantly higher rates relative to *S*. Typhi (STY). **E.** Scanning electron micrographs of STM-, or STY-infected ileum HIODEMs. STM infection is accompanied by cell rounding and barrier destruction, whereas STY infection maintains barrier function and showed bacterial association with the microvilli of enterocytes. Magnifications range from 7,000X to 14,000X, with 2 μm scale bars highlighted for each image. Please refer to Supplemental Figure S3 for additional images.

### Model validation with pathogenic *Escherichia coli*

Because the goal of the HIODEM system is to provide the most human-specific physiological model to analyze multiple enteric pathogens, validation of pathogenic *E. coli* infection would help support use of the model for a variety of bacteria. Since *Shigella* and *Salmonella* are both invasive pathogens, we decided to test the efficacy of our model on adherent, non-invasive pathogens while also reproducing similar analyses performed by another group (11). Enteropathogenic (EPEC strain 2348/69), enterohemorrhagic (EHEC O157:H7 strain 933), and enteroaggregative (EAEC strain 042) *E. coli* infect different locations of the intestine (*i.e*., ileum (34), colon (35), and colon (36), respectively). The ability of EPEC, EHEC, and EAEC to adhere or resist gentamicin was determined using organoids derived from the appropriate anatomical site specific to each pathogen in which the HIODEM monolayers were treated with DAPT only. Robust adherence for the three pathogens was detected in colon and ileum organoids; however, recoveries were minimal following gentamicin treatment (**Figure 6A**). To confirm these data, confocal immunofluorescence analysis (**Figure 6B**), SEM (**Figure 6C**), and TEM (**Figure 6D**) were performed. EPEC and EHEC adhered to the apical surface of the monolayers in tight association and indications of pedestal formation (37) were present, whereas EAEC adhered in an aggregative fashion (38). Interestingly, we found that the *E. coli* pathovars displayed a “hot-spot” infection pattern that was reproducible and consistent when imaged by various microscopy analyses. These results confirm the efficacy of the HIODEM model in recapitulating known phenotypes associated with pathogenic *E*. coli (11) and demonstrate the ability of the model to accommodate different pathogens for human cell infection analyses.

**Figure 6.**
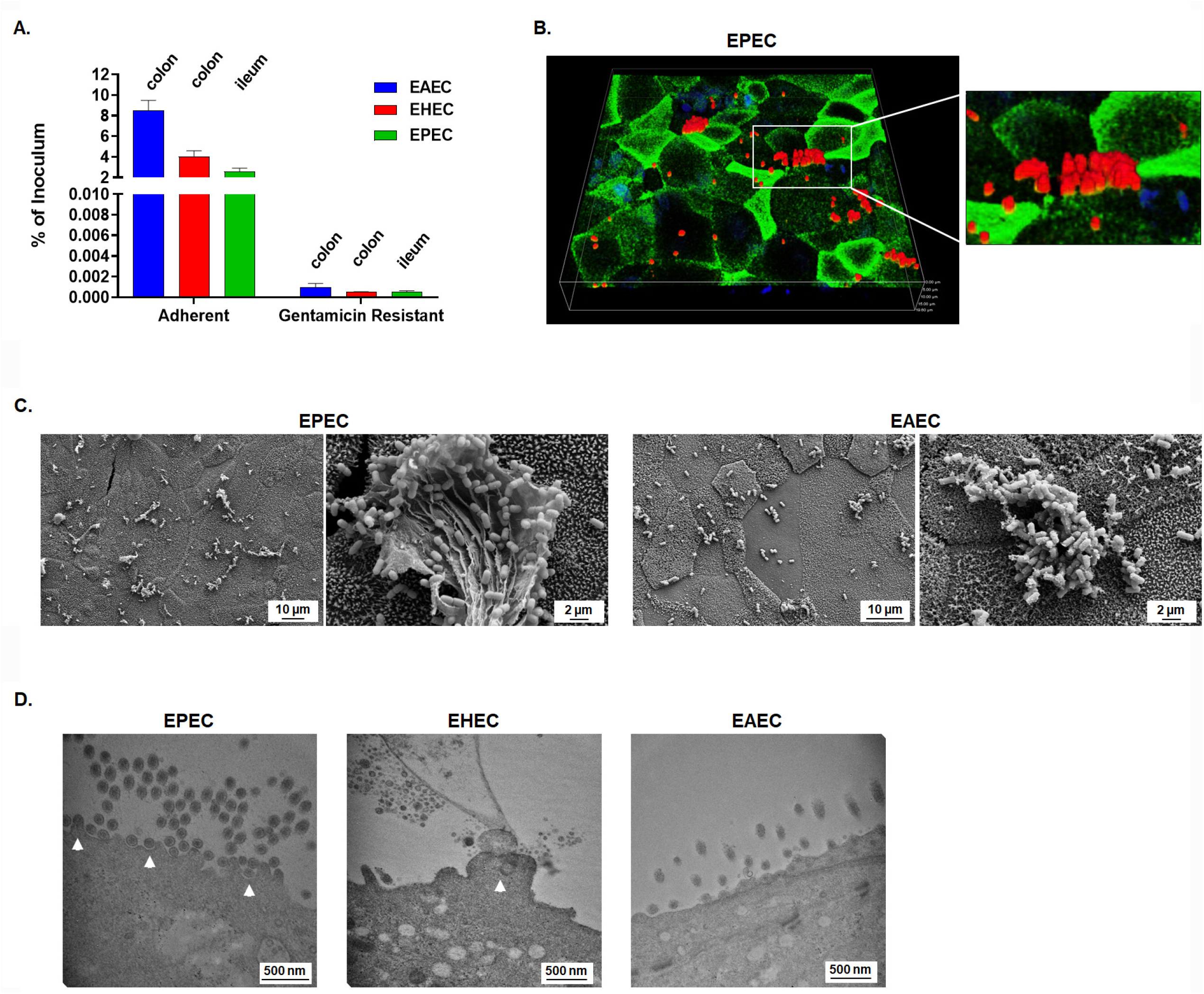
HIODEM supports infection with different *Escherichia coli* pathovars. **A.** Enteroaggregative (EAEC strain 042), enterohemorrhagic (EHEC 0157:H7 strain 933) and enteropathogenic (EPEC strain 2348/69) *E. coli* adhere to the surface of HIODEMs without significant invasion detected. Colon-derived HIODEMs was used for EHEC and EAEC, while ileum-derived HIODEMs were used for EPEC. **B.** Three dimensional (3D) confocal reconstructions of ileum HIODEM immunostained for actin (green), EPEC (red), and nuclei (blue) show recruitment of actin to the base of adherent bacteria in ileum-derived HIODEM. Magnification of the full image is 40X, with the height and width of the viewing plane at 141.3 μm and the depth at 19.6 μm. **C.** SEM of EPEC and EAEC infected monolayers demonstrate bacterial association with the host cells, aggregation of bacteria, and clusters of bacteria embedded in mucus. Magnifications range from 2,000X to 10,000X, with 10 μm and 2 μm scale bars, respectively. Ileum-derived HIODEM was used for EPEC and colon-derived HIODEM was used for EAEC. **D.** TEM images of infecting EPEC, EHEC and EAEC. Magnifications range from 50,000X to 60,000X with 500 nm scale bars indicated. White arrows indicate pedestal formation for EPEC and EHEC. Ileum- derived HIODEM was used for EPEC and colon-derived HIODEMs were used for EHEC and EAEC.

## Discussion

The use of human-derived intestinal organoids to study bacterial pathogenesis has recently increased (39-42). Prior to this advancement, most enteric pathogenesis studies were limited to immortalized cell lines and/or animal models. While traditional models have provided key insights into our understanding of enteric bacterial pathogens, successful vaccine development has been ineffective or limited (43-45). Animal models do not faithfully replicate the human GI tract (46-50) while immortalized cell lines have genetic abnormalities, dysregulated cell signaling pathways, and varied differentiation statuses depending on the cell line and the culturing conditions used (22, 51-53). Thus, there is a significant need to utilize human-specific infection models to improve our understanding of pathogenesis and to facilitate the development of efficacious vaccines against *Shigella, Salmonella*, and pathogenic *E. coli*. Intestinal-derived organoid models provide a pluripotent platform to study host-pathogen interactions, in which differentiation reagents ensure the presence of multiple cell types of the human epithelium while regional specificity can be retained by obtaining biopsies from different segments of the GI tract. Given the complexity of the methodology and reagents, we have provided a detailed protocol of our procedure (**Supplemental Methods**) to share with the research community and facilitate host-pathogen studies.

Organoid-derived epithelial monolayer models like the HIODEM have enabled infection analyses by providing a process by which pathogens can directly interact with the apical side of the epithelium (11, 24, 33, 54-58). While three-dimensional organoid systems are available (40, 59-63) and “apical-out” organoid systems have been developed (64), HIODEM and other 2-D systems offer easily accessible monolayers for pathogenesis studies in which apical and basolateral cytokine secretion profiles and changes in TEER can be examined in a standard transwell culture setting. Similar to traditional polarized models with Caco-2 and T84 cells (65-67), the polarized monolayer of HIODEM allows for a robust infection without need of centrifuging bacteria onto the cells, which is a common practice for infection analyses performed with standard, nonpolarized cell lines (68-70). Finally, the high-throughput nature of the multi-well plate formats enable multiple monolayers to be assayed simultaneously. Possible comparative analyses include various bacterial strains or mutants, infection conditions, ligand treatment, or even screening platforms for therapeutic candidates (23). The possibilities for therapeutic evaluation are amplified by the ability to develop organoids from donors with different genetic backgrounds, environmental exposure, age, disease status, or tissue site. These factors are captured and maintained in the organoid cultures (15, 71, 72), thereby increasing the potential of the model for patient stratification and precision medicine applications.

For our HIODEM system, media composition and inhibitors were carefully chosen to maintain the stem cell phenotype during the organoid culturing phase, initiate differentiation when seeded onto monolayers, and promote terminal differentiation in the last 48 hours of monolayer culture prior to use (see **Supplemental Methods**). Removal of the A-8301 inhibitor from the media when shifting from the organoid culture (1:1 + A-8301 + Y-27632) to the monolayer culture (1:1 + Y-27632) promotes cell attachment to the transwells (Dr. Senger’s observations). As the monolayer culturing progresses, TEER measurements monitor polarization of the models, which will vary depending on the tissue of origin. As noted in **Table 1**, our TEER values were consistent with published values for other cell-based models on transwell systems in which measurements are highest in the proximal intestine and decrease toward the colon. Passage of fluorescein isothiocyanate (FITC) dextran from the apical medium to the basolateral medium is another method to confirm the TEER readings and integrity of the barrier, as we have previously performed (15, 33, 71). Finally, transitioning the cells from the monolayer growth medium (apical and basolateral 1:1 + Y-27632) to terminal differentiation medium (apical cDMEM/F12 + DAPT, basolateral 1:1) prior to infection experiments promotes final differentiation since the factors that maintain stemness are significantly reduced and only present on the basolateral side to replicate *in vivo* environments (73). Thus, the removal of apical stem cell factors in conjunction with the addition of DAPT are critical signals that prompt the cells to differentiate and mature into a tissue-like model. It is important to note that each HIODEM model retains the cellular programming from the site of origin, *i.e*. ileum, cecum, or colon, despite the overall pluripotent state of the stem cells (11, 15, 71).

One of our goals was to develop a physiologically-relevant infection model for *Shigella*. Given the infection paradigm in which *Shigella* requires the presence of antigen-sampling M cells to access the basolateral pole for invasion of colonic epithelial cells (28), we sought to utilize a reagent that promotes the differentiation of M cells. We used RANKL based on previous studies (74, 75). The RANKL treatment resulted in significant induction of SPIB expression as detected by flow cytometry (**Figure 1**) and RT-qPCR (**Figure 2**), as well as increased appearance of the M cell phenotype upon microscopic examination of the monolayers (**Figure 3**). These results were consistent with previous analyses (74, 75). The differences in the levels of SPIB expression in each model following RANKL treatment likely reflect the retention of cellular programing from the original biopsy source, as noted above. Importantly, the expression of other cellular markers was not significantly altered with the use of RANKL (**Table 2** and **Supplemental Figure 1**). Thus, our results demonstrate that the cellular populations of the HIODEM models are capable of manipulation by ligand treatment to enable studies focused on specific cell types.

To examine the efficacy of the HIODEM models for pathogenesis, we evaluated *Shigella, Salmonella*, and pathogenic *E. coli* infection in models derived from the terminal ileum, cecum, and colon. Overall, we detected tissue tropism, pathogen targeting of specific cell types, and interesting infection dynamics by using models derived from the natural sites of infection. For *Shigella* (**Figure 4** and **Supplemental Figure S2**), adherence was highest in the cecum and colon, with the rates of both comparable to adherence rates with polarized T84 cells (26). Apical surface adherence was facilitated by *Shigella* adherence factors expressed in the *in* vivo-like culture conditions, and replicates previous analyses with both the colon and cecum models (23, 24, 26). Meanwhile, *Shigella* invasion was the highest in colon following treatment with RANKL to induce the presence of M cells. To confirm our data, we repeated analyses with another patient-derived colon model, and detected approximately a 10fold increase in recovery in which the DAPT + RANKL treatment increased *S. flexneri* colonic invasion relative to DAPT treatment (data not shown). Overall, both sets of invasion rates were reduced relative to rates seen in nonpolarized HeLa cells (69) or direct basolateral administration of polarized T84 cells (67, 76), but these models promote almost complete infection of the host cells and bypass important steps during the infection process. Since the HIODEM model is composed of various cell types, we only expect *Shigella* infection of the enterocytes. Our results agree with two recent publications using variations to the organoid-derived monolayer model that also verified the colonic tropism and the basolateral entry preference for *Shigella* invasion (54, 55). In this study, microscopy analyses indicated bacterial transit of M cells, which complements both previous confocal microscopy analyses indicating *Shigella* translocation of cecum-derived organoids following apical administration (23) and the invasion data rates obtained in the analyses described here. Furthermore, infection with the virulence plasmid-cured strain BS103 resulted in minimal recovery of the bacteria following gentamicin treatment, confirming that wild type 2457T recoveries in the presence of gentamicin were due to bacterial invasion of the model. In all, the bacterial culture conditions, apical administration of the inoculum, and the use of RANKL to promote M cell differentiation enables human-specific conditions to replicate the natural *Shigella* infection process in the laboratory setting.

In the literature, both *S*. Typhi and *S*. Typhimurium have been reported to infect the ileum (77-81), but result in differing pathologies: either systemic infection in the case *S*. Typhi or localized gastroenteritis in the case of *S*. Typhimurium. To explore this phenomenon in the HIODEM model, we analyzed infection of both serotypes in the ileum, cecum, and colon (**Figure 5** and **Supplemental Figure S3**). As with the *Shigella* analyses, we used bacterial culture conditions to induce virulence factor expression. Thus, *S*. Typhi was cultured with high-salt media and static growth to as previously described (33, 82) and the same protocol was used for *S*. Typhimurium to maintain similar growth conditions. The infection analyses demonstrated a surprising result that *S*. Typhi preferentially infected the cecum, while *S*. Typhimurium preferentially infected the ileum and caused more cellular destruction relative to *S*. Typhi. Furthermore, treatment of ileum-derived HIODEM monolayers with RANKL to induce M cell differentiation did not enhance *S*. Typhi invasion, which contrasts with *S*. Typhimurium observations (75) but is consistent with our previous analyses that *S*. Typhi invades human biopsies via the apical surface of enterocytes in which no bacterial associations with M cells were detected (33). Thus, the HIODEM model reproduced the serovar-specific differences that are expected given the different GI pathologies associated with each pathogen, while also identifying a unique cecum-specific infection pattern for *S*. Typhi. Reports of *S*. Typhi infection or damage to the cecum have been documented in the literature. Colonoscopic evaluations of patients with typhoid fever have found intestinal lesions in the terminal ileum in all patients tested, with additional lesions identified by the ileocecal valve and ascending colon (83). Furthermore, perforations of the ileocecum or lower gastrointestinal bleeding associated with the cecal artery have been reported in patients with *S*. Typhi infection (84, 85). Therefore, the HIODEM system offers opportunities to understand key differences between *S*. Typhi and *S*. Typhimurium, which may correlate with genetic differences of the pathogens (86) and/or epigenetic differences in humans.

Finally, the *E. coli* pathovar analyses validated the HIODEM system as a human-specific model that recapitulates expected infection patterns. Robust infection of the appropriate site of GI tract was observed with each pathovar, with comparable rates of adherence and minimal recoveries upon gentamicin to indicate a lack of invasion. The infection patterns and microscopic evaluations are in agreement with previous and recent analyses of organoid monolayer-based systems (11, 56, 57, 72), as well as with established literature regarding actin association and pedestal formation, or aggregative adherence patterns (37, 38). As with *Shigella* and *Salmonella* analyses, the future applications of the HIODEM model system with all *E. coli* pathotypes are broad and expected to provide key insights into human-specific pathology, including infection analyses in other segments of the GI tract for each pathovar.

In summary, we have provided reproducible infection analyses of six bacterial pathogens spanning three genera, in which unique observations have already been provided and infection data have been verified with electron microscopic analyses. This model and approach can be applied to additional enteric pathogens or bacteria representing the human microbiota. Coupled with *in vivo*-like bacterial culture conditions, the HIODEM model offers one of the most human-specific infection analyses that can be performed in the laboratory setting to further our understanding of host-microbe interactions and hopefully help lead to the discovery of novel vaccines and therapeutics.

## Methods and Materials

### Human Subjects Research, IRB Approval, and Biopsy Collection

Human sample collection was approved by Institutional Review Board (IRB) protocols 2014P002001 and 2015P001908 of the Massachusetts General Hospital, Boston, MA. Donor tissue was obtained from consenting patients undergoing medically required colonoscopies or surgical resections, as determined by a licensed physician. All subjects provided written informed consent for samples to be used for research purposes.

### Organoid Culture and Monolayer Generation

The protocol has been adapted from previous publications (11, 15, 33). Please refer to the **Supplemental Methods** for step-by-step instructions. Briefly, stem cells derived from donor biopsies were maintained in Matrigel culture in a 1:1 mixture of intestinal stem cell media (ISC) + L-WRN conditioned media containing the inhibitors Y-27632 and A- 83-01. Cells were seeded at a density of 15,000 cells per Matrigel dome and grown for 7 days in culture. Spheres were trypsinized into a single-cell state, seeded onto polyethylene terephthalate (PET) membrane transwell inserts with a 0.4-μm pore size at 1.0 × 10^6^ cells/ml, and incubated in the 1:1 stem cell medium-L-WRN medium at 37°C with 5% CO_2_. The culture medium was changed every other day until the cultures reached confluence, as determined by transepithelial electrical resistance (TEER) monitoring and microscopic observation. Cells grew and matured for 7-10 days, at which time the apical and basolateral media were changed and differentiation reagents were applied for either 24 or 48 hours depending on the experiment. Differentiation reagents included 5 μM γ-secretase inhibitor IX (DAPT; Calbiochem) application to the apical surface, which was also combined with 100 or 500 ng/ml of the receptor activator of the NF-κB ligand (RANKL; Peprotech) application to the basolateral media where indicated.

### Infection analyses

Please refer to the detailed protocols in the **Supplemental Methods**.

### Transepithelial electrical resistance (TEER)

To assess paracellular permeability, transwell inserts were monitored using a TEER apparatus (World Precision Instruments, Sarasota, FL) per manufacturer’s instructions (15, 33, 66).

### Reverse transcription quantitative PCR (RT-qPCR) analysis

RNA was extracted from monolayers using Trizol (ThermoFisher) and Direct-Zol (Zymo) RNA extraction kit. RNA was treated with on-column DNase. RNA was converted to cDNA using Thermo Scientific Maxima First Strand cDNA Synthesis Kit. Gene expression quantitation was determined using Sybr Green (Perfecta) and The CFX96 real-time polymerase chain reaction (PCR) detection system (Qiagen, Venio, NL) was used for gene expression analysis. The relative threshold cycle (ΔΔCT) method was used for assessing gene expression relative to the 18S housekeeping reference gene (15, 33). Identification of cell type was based on the following genes: *LGR5* (87-89) for stem cells; *ESE1* (19) and *SIM* (sucrose isomaltase) (90) for enterocytes; *KLF4* (20) and *MUC2* (91) for goblet cells; *SPIB* for M cells (17); *CGA* (92) and *NEUROD* (93) for enteroendocrine cells; *OCLDN* (occludin) (94, 95) for barrier formation; and *SOX9* (96, 97), *SPDEF* (96, 98), and *LYZ* (lysozyme) (99, 100) for Paneth cells.

### Flow Cytometry

After differentiation, large transwells (12-well plates) were trypsinized for 10 minutes. Cells were subsequently resuspended in cDMEM and kept on ice until staining. Cells were fixed, permeabilized, and stained with flow cytometry fixation and permeabilization buffer kit as directed by the manufacturer. Briefly cells were washed twice with 1X PBS (Gibco), resuspended in 500 μl of Flow Cytometry Fixation buffer (R&D), and incubated for 10 minutes at room temperature. Following fixation, cells were permeabilized with 200 μl of Flow Cytometry Permeabilization/Wash buffer (R&D), stained with anti-human ESE1 (Abcam), anti-human KLF4 APC-conjugated (R&D System) and anti-human SPIB (Invitrogen) antibodies by incubating for 45 minutes at 4°C. Afterwards, the excess antibodies were removed by washing the cells with Flow Cytometry Permeabilization/Wash buffer and stained with secondary antibodies anti-rabbit IgG1 FITC (Abcam) and anti-mouse IgG2 PerCp (BD Bioscience) for 20 minutes at 4°C. Samples were washed again with Flow Cytometry Permeabilization/Wash buffer, fixed in 1% paraformaldehyde in 1X PBS (Gibco), and acquired with BD FACSCalibur flow cytometer. Analysis was performed with BD bioscience software. Enterocytes (ESE1^+^), M cells (SPIB^+^), and Goblet cells (KLF4^+^) were gated among live cells based on FSC and SSC parameters.

### Electron microscopy

For transmission electron microscopy (TEM) analysis, samples were fixed in 2% paraformaldehyde/2.5% glutaraldehyde in 0.1 M Sodium Cacodylate followed by mounting on grids and imaged using a transmission electron microscope (JEOL, Peabody, MA). For scanning electron microscopy (SEM) analysis, HIODEM monolayers were fixed in 0.5X Karnovsky fixative (Newcomer Supply) and subsequently stored in 1X PBS at 4°C. All sample processing occurred at the Massachusetts Eye and Ear Infirmary core facility. All SEM imaging was performed at the Harvard University Center for Nanoscale Systems (CNS) using a FESEM Supra55VP microscope.

### Cytokine Analysis

Quantification of secreted interleukin 8 (IL-8) was conducted using R&D Systems Human CXCL8/IL-8 DuoSet ELISA per manufacturer’s instructions (101).

### Lactate dehydrogenase (LDH) assay

Apical supernatants were assessed for LDH release using Promega Cytox Kit (Promega, Madison, WI) according to manufacturer’s instructions (33).

### Immunostaining

For immunostaining, monolayers were fixed in 4% paraformaldehyde at room temperature for 15 minutes followed by storage in 70% ethanol at 4°C until paraffin embedding (33). Embedding and sectioning were performed by the Specialized Histopathology Core of Massachusetts General Hospital. Prior to staining, sections were deparaffinized using xylene with gradual rehydration in decreasing concentrations of ethanol. Sections were blocked using 0.4% goat and donkey serum in 0.04% Triton X-100 in PBS. Sections were stained using the antibodies against actin (Cell Signaling Technologies, 3700S), Mucin 2 (Santa Cruz Technologies, sc-13312 (33)), *Salmonella* (Bio-rad, 8209-4006) and *E. coli* (Abcam, ab137967, kind gift of Deepak VK Kumar). Fluorescently conjugated secondary monoclonal antibodies (Alexa Fluor 488 and 555 conjugated antibody series against mouse, rabbit, or goat from Life Technologies) were used for detection. Nuclei were counterstained with 6- diamidino-2-phenylindole (DAPI). Samples were imaged using a Nikon A1SiR confocal microscope.

## Acknowledgements

The authors gratefully acknowledge Dr. Francis Colizzo, Dr. James Michael Richter, and Dr. Barbara Nath for collection of the samples. Without their technical expertise, these studies would not have been possible. We also thank Dr. Bobby Cherayil and Dr. Brian Hurly for bacterial strains used in this study, Ms. Diane Capen for her expertise and skill in preparing the samples for TEM analysis, Ms. Ann Tisdale for her expertise in preparing the SEM samples, and Dr. Tim Cavanaugh and the Center for Nanoscale Studies at Harvard University for use of the SEM. We would also like to thank Dr. M. Rosaria Fiorentino, members of the Fasano, Fiorentino, and Faherty laboratories, Dr. Beth McCormick at the University of Massachusetts Medical School, as well as the members of the University of Maryland Cooperative Center on Human Immunology (CCHI) for their thoughtful feedback and discussions during the project.

This work was supported by the National Institute of Allergy and Infectious Diseases grants K22 AI104755 (CSF), and R01-AI036525 (MBS), NIH U19-AI082655 Cooperative Center on Human Immunology (MBS and AF), and DHHS U19-AI109776 (Center of Excellence for Translational Research, CETR, MBS). The TEM core is supported by National Institute of Neurological Disorders and Stroke P30NS045776. Support for the Philly Dake Electron Microscope Facility was provided by the National Institutes of Health grant 1S10RR023594S10 and by funds from the Dake Family Foundation. The Dana-Farber/Harvard Cancer Center Specialized Histopathology Core is supported, in part, by an NCI Cancer Center Support Grant # NIH 5 P30 CA06516. The funders had no role in study design, data collection and analysis, decision to publish, or preparation of the manuscript. Dr. Senger discloses the following conflicts of interest: PFE, BABA, and KLDO stock holdings.

## Supplemental Files

**Supplemental Figure S1. Additional flow cytometry data.** The percentage of each cell type (enterocytes (ESE1), goblet cells (KLF4), and M cells (SPIB)) detected by flow cytometry for the terminal ileum, cecum, and colon. The data represent the average percentage +/- SEM of each cell population at 24- and 48-hours following differentiation, in which treatments included apical DAPT (black bars), or a combination of apical DAPT with basolateral physiological 100 ng/ml RANKL treatment (red bars) and supraphysiological 500ng/mL (blue bars). Statistical significance was determined with a one-way ANOVA (paired Friedman test) for the samples treated with RANKL versus the DAPT only treatment of the matched originating tissue (* = p<0.05). Please note that the SPIB data corresponds to the data presented in Figure 1.

**Supplemental Figure S2. Additional analysis of *Shigella flexneri* HIODEM infection.**

**A.** Colon-derived HIODEM monolayers were treated with DAPT + RANKL. Uninfected monolayers (media) or infected with *S. flexneri* demonstrate significant secretion of IL-8 after infection. Statistical significance was determined by the Student’s t-test, * = p<0.05.

**B.** No significant increase in cell death as evaluated by LDH-release was detected following infection of colon-derived HIODEM monolayers treated with DAPT + RANKL.

**Supplemental Figure S3. Additional microscopy analysis of *Salmonella* HIODEM infection.**

**A.** Confocal immunofluorescence analysis of *S*. Typhi HIODEM infection. Images of control (left) or *S*.

Typhi infection (right) of ileum-derived HIDOEM. Samples were stained with DAPI to visualize cellular DNA (blue), α-Muc2 to identify the mucus barrier secreted from goblet cells (green) and *α-Salmonella* to visualize the infecting population (red). Colocalization of *Salmonella* and DAPI results in the bacteria appearing purple.

**B.** Scanning electron micrographs of STM-, or STY-infected ileum HIODEMs. As demonstrated in Figure 5, STM infection led to significant destruction of the barrier, while STY infection did not affect the barrier and the bacteria associated with microvilli on the apical surface of the enterocytes. 2 μm scale bars are indicated.

**Supplemental Methods.** Expanded methodology for culturing organoids and plating, differentiating, and infecting the monolayers with *Shigella, Salmonella*, or the different *E. coli* pathovars.

